# Lack of susceptibility of poultry to SARS-CoV-2 and MERS-CoV

**DOI:** 10.1101/2020.06.16.154658

**Authors:** David L. Suarez, Mary J. Pantin-Jackwood, David E. Swayne, Scott A. Lee, Suzanne M. DeBlois, Erica Spackman

**Author notes:** Corresponding author: **Address for Correspondence:** Erica Spackman, US National Poultry Research Center, USDA-Agricultural Research Service, 934 Station Rd., Athens, GA 30605, USA.

## Abstract

Chickens, turkeys, ducks, quail and geese were challenged with SARS-CoV-2 or MERS-CoV. No disease was observed, no virus replication was detected, and antibodies were not detected in serum. Neither virus replicated in embryonating chicken’s eggs. Poultry are unlikely to serve a role in the maintenance of either virus.

---

Coronaviruses (CoV) of animals periodically transmit to humans *(1)* as recently occurred with SARS-CoV-2. SARS-CoV-2 was first recognized in December of 2019 with cases of atypical pneumonia in hospitalized patients in Wuhan, China. The virus is a novel beta-coronavirus, related to the now eradicated, severe acute respiratory syndrome (SARS) CoV from 2003 with which it has 82% identity across the genome *(2)*. SARS-CoV-2 is highly transmissible among humans and particularly virulent for elderly individuals and those with certain underlying health conditions. Multiple studies have examined the susceptibility of domestic animals to SARS-CoV-2 to establish the risk of zoonotic transmission and two studies have shown chickens and Pekin ducks were not susceptible to infection *(3, 4)*.

Middle East Respiratory Syndrome coronavirus (MERS-CoV), another coronavirus of high concern associated with zoonotic infection, was first detected in patients with severe acute lower respiratory tract disease in Saudi Arabia in 2012. MERS-CoV causes lower respiratory disease similar to the SARS-CoVs *(5)*. Unlike SARS-CoV-2, MERS-CoV transmits poorly to humans and does not exhibit sustained human-to-human transmission; however, it has a high case fatality rate of around 30%. Although the MERS-CoV case count is low, human cases continue to be reported, therefore there is a possibility for the virus to adapt to humans.

Based on sequence similarity, the closest relatives of SARS-CoV-2 and MERS-CoV are believed to be bat beta-coronaviruses *(6)*, but because of the amount of sequence difference between human and bat isolates an intermediary host likely exists. For MERS-CoV, dromedary camels appear to be the primary natural reservoir of infection to humans, but other domestic animals seem to be susceptible to infection *(7, 8)*. There is only a single study of MERS-CoV in chickens that looked for antibodies, but all samples were negative *(9)*.

Because poultry are so widespread and have close and extended contact with humans, and other mammals in many production systems, including live animal markets, susceptibility were conducted with SARS-CoV-2 and MERS-CoV in five common poultry species. Additionally, embryonating chicken eggs (ECE) have been utilized for the isolation and as a laboratory host system, including use in vaccine production, for diverse avian and mammalian viruses. Therefore, ECE were tested for their ability to support the replication of both viruses.

Five poultry species were examined: chickens (*Gallus gallus domesticus*), turkeys (*Meleagris gallopavo*), Pekin ducks (*Anas platyrhinchos domesticus*), Japanese quail (*Coturnix japonica*) and White Chinese geese (*Anser cygnoides*). All procedures involving animals were reviewed and approved by the US National Poultry Research Center Institutional Animal Care and Use Committee, and the viruses were used under the approval of the Institutional Biosafety Committee.

To evaluate their susceptibility to these viruses, 10 birds of each species were challenged with either the USA-WA1/2020 isolate of SARS-CoV-2 (BEI NR-58221) or the Florida/USA-2_SaudiArabia_2014 isolate of MERS-CoV (BEI NR-50415), which were both obtained from the Biodefense and Emerging Infections Research Resources Repository (BEI Resources), National Institute of Allergy and Infectious Diseases, National Institutes of Health (full details of all methods are provided in the technical appendix). Oro-pharyngeal and cloacal swabs were collected from all birds at 2, 4, and 7 days post challenge (DPC) and were tested for virus by real-time RT-PCR. At 14 DPC sera were collected from the birds and were tested for antibody to the challenge virus by microneutralization.

Clinical signs were not observed at any time in any species, and virus was not detected in any swab material. Antibodies were not detected in serum from any birds at 14 days post challenge. These results suggest that neither virus replicated in any of the avian species evaluated or replicated at a level that was too low to be detected.

ECE were tested for their ability to support SARS-CoV-2 or MERS-CoV replication after inoculation with any of the three most common routes: yolk sac, chorio-allantoic sac and chorio-allantoic membrane (see technical appendix for details). Yolk, allantoic fluid/albumin, and embryo tissues were collected from inoculated eggs and tested for viral replication by attempting virus isolation in Vero cells from the egg material after each of 2 ECE passages. Neither virus was recovered in Vero cells from the inoculated ECEs and lesions were not observed in any of the embryos inoculated with SARS-CoV-2 or MERS-CoV.

Identifying potential reservoir hosts of the novel coronaviruses is critical to controlling exposure and subsequent infection, as well as to preserving a safe and consistent food supply. None of the avian species evaluated here, nor ECE appeared to support replication of either virus. Therefore, poultry are unlikely to serve a role in the maintenance or transmission of either SARS-CoV-2 or MERS-CoV. Further, ECE are not a viable laboratory host system.

## Supporting information

Technical Appendix

## Acknowledgments

The authors gratefully acknowledge Jesse Gallagher, Melinda Vonkungthong, Anne Hurley-Bacon, Jasmina Luczo, James Doster, and Charles Foley for technical assistance with this work.

The following reagent was deposited by the Centers for Disease Control and Prevention and obtained through BEI Resources, NIAID, NIH: SARS-Related Coronavirus 2, Isolate USA-WA1/2020, NR-52281. The following reagent was obtained through BEI Resources, NIAID, NIH: Middle East Respiratory Syndrome Coronavirus, Florida/USA-2_Saudi Arabia_2014, NR-50415. Vero African Green Monkey Kidney Cells (ATCC® CCL-81™), FR-243, was obtained through the International Reagent Resource, Influenza Division, WHO Collaborating Center for Surveillance, Epidemiology and Control of Influenza, Centers for Disease Control and Prevention, Atlanta, GA, USA.

This work was supported by USDA-Agricultural Research Service project #6040-32000-066-00-D. Mention of trade names or commercial products in this manuscript is solely for the purpose of providing specific information and does not imply recommendation or endorsement by the U.S. Government. USDA is an equal opportunity provider and employer. Paragraph listing persons or organizations you wish to acknowledge.

## Author Bio (first author only, unless there are only 2 authors)

Dr. David Suarez is the Research Leader for the Exotic and Emerging Avian Viral Disease Research Unit of the Agricultural Research Service, USDA. His primary research interests are in the understanding and control of avian influenza and Newcastle disease viruses in poultry and other emerging viral diseases that threaten the poultry industry.

