## Supplementary material for "Lack of susceptibility of poultry to SARS-CoV-2 and MERS-CoV": Technical Appendix

### DETAILED METHODS

**Viruses**. The USA-WA1/2020 (BEI NR-58221) *(1)* isolate of SARS-CoV-2 and the Florida/USA-2_SaudiArabia_2014 isolate of MERS-CoV (BEI NR-50415) *(2)* were both obtained from Biodefense and Emerging Infections Research Resources Repository, National Institute of Allergy and Infectious Diseases, National Institutes of Health. Both viruses were propagated and titrated in CCL-81 Vero cells (International Reagent Resource FR-243). SARS-CoV-2 was utilized at 5 total passages in Vero cells, and MERS-CoV was utilized at 6 total passages in Vero cells. Viruses were used under the approval of the US National Poultry Research Center Institutional Biosafety Committee.

**Evaluation of virus replication in avian species.** Five poultry species were selected because of their prevalence worldwide: chickens (*Gallus gallus domesticus*), turkeys (*Meleagris gallopavo*), Pekin ducks (*Anas platyrhinchos domesticus*), Japanese quail (*Coturnix japonica*) and in wet markets in China: chickens, Pekin ducks, quail, and Chinese domestic geese (*Anser cygnoides*). Chickens and turkeys were obtained from in-house specific pathogen free (SPF) flocks. Ducks, geese and quail were obtained from a commercial hatchery. All procedures involving animals were reviewed and approved by the US National Poultry Research Center Institutional Animal Care and Use Committee.

Each bird was individually tagged for identification. For each species 10 birds were challenged with each virus and three birds were not inoculated to serve as age matched controls.

Blood was collected from all birds immediately prior to infection and was tested by microneutralization for antibodies to the appropriate challenge virus. Chickens, turkeys and quail were challenged at 4 weeks of age, ducks and geese were challenged at 2 weeks of age (Table S1). Chickens, turkeys and quail were challenged with 5.4 $\log_{10}$ 50% tissue culture infectious doses ($TCID_{50}$) of SARS-CoV-2 in 0.1mL or 5.2$\log_{10}$ $TCID_{50}$ of MERS-CoV in 0.1mL by the intra-choanal route. Ducks and geese were challenged with 6.0 $\log_{10}$ $TCID_{50}$ of SARS-CoV-2 or 5.5 $\log_{10}$ $TCID_{50}$ of MERS-CoV, each in 0.1mL by the intra-choanal route. Birds were observed a minimum of daily for clinical signs.

Oro-pharyngeal (OP) and cloacal (CL) swabs were collected from all challenged birds at 2, 4, and 7 days post challenge (DPC) and were tested for virus by real-time RT-PCR. The rRT-PCR was run with the 2, 4 and 7DPC samples immediately after the 7DPC samples were collected. Because they were negative, it was determined that it was not necessary to test any later time points.

Because there was no evidence of infection, no clinical signs and no virus was excreted by the respiratory tract or intestinal tracts, lesions were not expected to have developed, therefore no birds were necropsied during the course of the study.

At 14 DPC blood was collected from all surviving birds and the sera were tested by microneutralization to evaluate whether there was an antibody response to the challenge virus.

**Table S1.** Age at challenge and dose for each virus by species ($TCID_{50}$ = 50% tissue culture infectious dose)

| Species | Age at challenge | Titer of challenge with SARS-CoV-2 ($log_{10}$ $TCID_{50}$/bird) | Titer of challenge with MERS-CoV ($log_{10}$ $TCID_{50}$/bird) |
|---|---|---|---|
| Chickens (*Gallus gallus domesticus*) | 4 weeks | 5.4 | 5.2 |
| Turkeys (*Meleagris gallopavo*) | 4 weeks | 5.4 | 5.2 |
| Japanese quail (*Coturnix japonica*) | 4 weeks | 5.4 | 5.2 |
| Pekin ducks (*Anas platyrhinchos)* | 2 weeks | 6.0 | 5.5 |
| Chinese domestic geese (*Anser cygnoides*) | 2 weeks | 6.0 | 5.5 |

**Replication in embryonating chicken's eggs.** Embryonating chicken eggs (ECE) were evaluated for their ability to support replication of SARS-CoV-2 and MERS-CoV. Procedures were identical for both viruses. Five ECE were inoculated with $10^{6.5}$ $TCID_{50}$ in 0.2mL for each of the 3 most common routes of inoculation: yolk sac (YS), chorio-allantoic sac (CAS), and chorio-allantoic membrane (CAM). Established inoculation procedures for each route were utilized *(3)*. Eggs were candled daily for viability.

Samples were collected from the inoculated eggs when the embryo was found to be non-viable or at the end of the incubation period. Yolk, allantoic fluid/albumin, embryo tissue (2-3 grams of viscera and thigh muscle) were collected from YS inoculated eggs. Allantoic fluid/albumin, embryo tissues were collected from CAS inoculated eggs, and allantoic fluid/albumin, embryo tissues and egg membrane were collected from CAM inoculated eggs. During sample collection the embryos were dissected to observe lesions. Age matched non-inoculated ECE served as controls.

CAM and embryo tissues were homogenized in PBS with glass beads in a FastPrep 24 (MP Biomedical LLC, Santa Anna, CA) then was centrifuged at 17Kxg for 10 minutes and the supernatant was used for the second passage and for RNA extraction for subsequent testing by rRT-PCR. Allantoic fluid/albumin was used directly for the second passage and RNA extraction. To complete the second passage all sample material from the 5 eggs of same inoculation route were pooled. The material was then inoculated identically to the first passage. Material from both passages was tested by inoculation into Vero cells in triplicate for fluid and embryo material from each inoculation route as described above to test for the presence of virus.

**Microneutralization.** Virus microneutralization with serum from each species was conducted with both SARS-CoV-2 and MERS-CoV in CCL-81 Vero cells as described by Algaissi and Hashem *(4)*, with the modification that the dilutions of serum tested were 1:4, 1:8, 1:16 and 1:32. Also, the antibody treated virus was added when the cells were plated. Titers greater than 1:8 were considered positive. Positive control antibodies were commercially available monoclonal antibodies to the S2 region of the spike protein: SARS-CoV-2 used at 12.5μg/mL (MP Bio, Irvine, CA), and MERS-CoV used at 20μg/mL (EastCoast Bio, North Berwick, ME).

**RNA extraction and quantitative real-time RT-PCR.** RNA was extracted from OP and CL swab material with the Ambion Magmax kit (ThermoFisher, Waltham, MA) as described previously *(5)*. The rRT-PCR primers, probe and cycling conditions for SARS-CoV-2 for the N1 primer and probe set from the US Centers for Disease Control was utilized *(6)*. The N3 primers, probe and conditions reported by Lu *et al*. which target the nucleoprotein gene was used for MERS-CoV detection *(7)*. The AgPath ID one-step RT-PCR kit was used and the RT step of the reaction conditions was modified to accommodate the recommended kit conditions (PCR conditions recommended for each primer and probe set from the original protocols were used). A standard curve of RNA titrated virus was run in duplicate with each run of rRT-PCR to estimate titer equivalents of virus present in samples.

**Reagent Acknowledgements**

The following reagent was deposited by the Centers for Disease Control and Prevention and obtained through BEI Resources, NIAID, NIH: SARS-Related Coronavirus 2, Isolate USA-WA1/2020, NR-52281. The following reagent was obtained through BEI Resources, NIAID, NIH: Middle East Respiratory Syndrome Coronavirus, Florida/USA-2_Saudi Arabia_2014, NR-50415. Vero African Green Monkey Kidney Cells (ATCC® CCL-81™), FR-243, was obtained through the International Reagent Resource, Influenza Division, WHO Collaborating Center for Surveillance, Epidemiology and Control of Influenza, Centers for Disease Control and Prevention, Atlanta, GA, USA.
